# The social conditioned place preference assay is sensitive to drug dose, training regimen, and social context

**DOI:** 10.64898/2026.11.09.765097

**Authors:** Makenzie Wilkinson, Gül Dölen

## Abstract

Beginning in the 1960s, researchers have understood that one of the unique properties of psychedelic drugs is that their effects are modified by the context in which they are taken (also known as their ‘set’ and ‘setting’ dependence). To date, 7 different studies have shown that critical period reopening is a shared property across psychedelic drugs, including ketamine, MDMA, psilocybin, LSD, and ibogaine. Moreover, the context dependence of psychedelics’ effects has been recapitulated for both MDMA and ketamine. Recently, Lu et al., *Nature Neuroscience* 2026 (PMID: 42972049) have claimed that psilocybin does not reopen the social reward learning critical period, which the authors interpret as a failure to replicate previously published results. Here, we re-analyzed results across three published datasets (Nardou et al., *Nature* 2019; Nardou et al., *Nature* 2023; Lu et al., *Nature Neuroscience* 2026) in order to compare the impact of dose, training regimen, and social context on the social conditioned place preference (social CPP) assay. Our analysis reveals that methodological modifications likely account for Lu and colleagues’ inability to reopen the social reward learning critical period with psilocybin. Nevertheless, because their results recapitulate previous findings that psychedelics’ effects are context dependent, they support rather than challenge the growing literature demonstrating that psychedelics reopen critical periods.

## Main

Lu and colleagues^1^, set out to test the reproducibility and reliability of previously published behavioral studies of psychedelics in mice, by utilizing uniform protocols repeated across 5 labs. Although the authors admit that they modified many of these protocols from the originally published versions, they argue that truly robust findings and generalizable effects should be significant even across slight variations in experimental conditions. Based on the results of their studies, the authors claim that only anxiogenic effects of psilocybin and psilocybin-mediated attenuation of fear expression are consistent across the consortium labs^1^ and conclude that their findings point to limitations in the robustness and reproducibility of other assays.

Focusing on the studies reported in Figure 7 of Lu et al.^1^, which were ostensibly designed to replicate findings from Nardou et al., Nature 2019 and 2023^2,3^, alterations to the social conditioned place preference assay (social CPP) were substantial and may have impacted the outcome of their study. Indeed, previous studies using the social CPP protocol have shown that this assay is sensitive to variations in a number of parameters including: odorants, bedding cues, training schedule, sleep, nesting, stress, age, background strain, and the rearing history of the animals (reviewed in Wilkinson and Dolen, *JoVE* 2026)^4^. Therefore, here, we re-examined the conclusions drawn by Lu et al., by making side by side comparisons of protocol modifications, including dose, training, and social context, and reanalyzed results across three published datasets (Nardou et al., Nature 2019; Nardou et al., Nature 2023; Lu et al., Nature Neuroscience 2026).

## Results

### Dose significantly impacts the subjective experience of psychedelic drugs

Compared to our studies (0.3 mg/kg) (**Figure 1a,b**), and indeed most other studies in the field, Lu and colleagues used an unusually high dose of psilocybin (2 mg/kg) (**Figure 1c,d**). Theoretically, 2 mg/kg in mice would be the equivalent of double to ten times the doses used in human clinical trials (even when correcting for species specific differences in 5-HT2A receptors)^5^. Thus, it is not surprising that one of the only consistent findings reported by Lu and colleagues is anxiety-like behavior in response to acute psilocybin. Significantly, this difference in dose could partially account for the failure to reopen the social reward learning critical period (**Figure 1d**; psilocybin 2 mg/kg; n = 106, *t*_(105)_ = 1.196, *P =* 0.2343), since it would substantially impact the ability of the animals to engage in a social context during the psilocybin trip (as described further below). Moreover, previous studies have shown that psilocybin’s ability to reopen the social reward learning critical period is dose dependent^3^, with 0.3 mg/kg being the lowest dose that induced this effect (**Figure 1b**; psilocybin 0.3 mg/kg: n = 15, *t*_(14)_ = -3.741, *P =* 0.0002). Given that both MDMA and ketamine have inverted U-shaped dose response curves, whereby higher doses fail to reopen the social reward learning critical period^3^, the results from Lu et al., raise the possibility that psilocybin mediated critical period reopening shares a similar dose response profile. Nevertheless, given other protocol differences described below, such conclusions are necessarily provisional.

**Figure 1.**
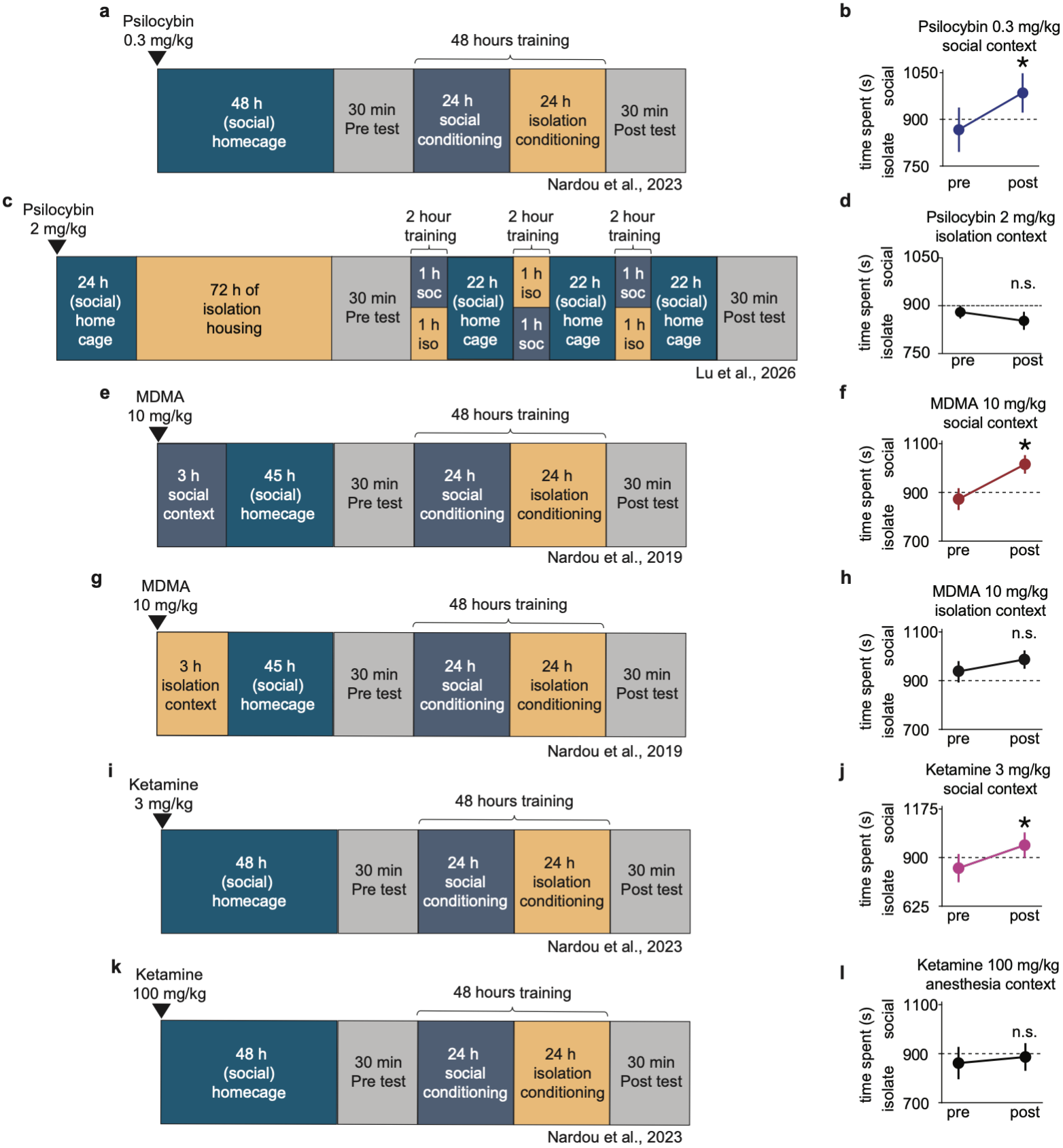
Significant alterations to social CPP protocol: dose, training, and context (a, c, e, g, i, k) Experimental time course of i.p. psychedelic pretreatment in sCPP highlighting differences between the original protocol used by Nardou et al., in 2023 (**a, i, k**), and 2019 (**e, g**) and the “simplified” protocol used in Lu et al., 2026 (**c**). (**b, d, f, h, j, l**) Average responses of adult mice comparing time spent on side conditioned to the social cue in the pre-vs post-test. Statistically significant differences between pre- and post-test indicate reopening of the social reward learning critical period. Comparisons revealed that (**b**) psilocybin (0.3 mg/ kg; n = 15, *t*_(14)_ = -3.741, P = 0.0002), (**f)** MDMA (10 mg/kg; n = 26, t_(25)_ = -4.116, *P <* 0.0004) and (**j**) ketamine (3 mg/kg; *n* = 18, *t*_(17)_ = -3.826, *P <* 0.002) reopened the critical period when administered in a social setting. This feature was not shared with (**d**) psilocybin (2 mg/kg; n =106, *t*_(105)_ = 1.196, *P =* 0.2343) or (**g**) MDMA (10 mg/kg; n = 29, *t*_(28)_ = -1.97, *P =* 0.059) when paired with an isolation setting, or (**l**) ketamine (100 mg/kg; *n* = 20, *t*_(19)_ = -0.89452, *P =* 0.382) under anesthesia. Data from Lu et al., 2026 (**c, d**) were converted from percentages into raw numbers to run an analogous analysis (two-tailed paired *t-*test). We observed the same result of no statistical difference between pre- and post-test time on the social bedding but disagree regarding the interpretation. All data are presented as mean ± s.e.m.

### Adjustments to sCPP training schedule make learning significantly more difficult

The authors substantially reduced the number of hours of training given to the animals (6 hours) compared to previous studies (48 hours). **Figure 1a** describes the sCPP protocol utilized by Nardou et al., 2019 and 2023, which consists of two back-to-back 24-hour training sessions on novel bedding cues trained to either the social (group housed with cage mates) or the isolation (singly housed) condition. The shortened training regimen used in Lu et al., 2026 is outlined in **Figure 1c**. Two-hour long training sessions (1 h social, 1 h isolation) were used across three consecutive days, each followed by removal from the training environment for the remaining 22 hours of each day. Although the authors claim that their protocol alteration is a “simplification” of the social CPP assay, this is hardly self-evident since the amended protocol takes 7 days (instead of 4) and has a complicated schedule involving 13 training/testing arena exchanges (as opposed to 5 in the original), each of which adds unnecessary handling stress to the protocol. These modifications are particularly dubious given that the last author has two prior publications^6,7^ where the protocol used for measuring social CPP was identical to the papers they are attempting to replicate here^2,3^. Furthermore, although these alterations did not interfere with their ability to reproduce our finding that social reward learning is governed by a critical period^2^, these shortened training regimens are substantially less robust^4,8^ which likely necessitated the introduction of other elements (e.g. social isolation, see below) to the protocol.

### Psychedelic effects are modified by the social context in which they are delivered

Lu and colleagues introduced 3 days of social isolation between the psychedelic drug dose and the measurement of the social reward learning critical period, substantially altering the context of the psilocybin experience (**Figure 1c**). Importantly, previous work has shown that the critical period open state induced by psilocybin (as in **Figure 1a**) persists for 2 weeks following drug administration^3^ during which time, the subjects would be sensitive to the social experience (or lack thereof). Consequently, this amendment to the protocol renders their experiment more similar to our previous control studies. Specifically, whereas MDMA given in a social context (**Figure 1e**) reopens the social reward learning critical period (**Figure 1f**; MDMA 10 mg/kg: *n* = 26, *t*_(25)_ = -4.116, *P <* 0.0004), MDMA delivered in a social isolation context (**Figure 1g**) fails to reopen the social reward learning critical period (**Figure 1h;** MDMA 10 mg/kg: *n* = 29, *t*_(28)_ = -1.97, *P =* 0.059)^2^. We have also previously shown that whereas ketamine given at low psychedelic doses (**Figure 1i**) reopens the social reward learning critical period (**Figure 1j;** ketamine 3 mg/kg: *n* = 18, *t*_(17)_ = -3.826, *P <* 0.002), higher anesthetic doses (**Figure 1k**) of ketamine fail to do so (**Figure 1l;** ketamine 100 mg/kg: *n* = 20, *t*_(19)_ = -0.89452, *P =* 0.382), presumably because the animals do not consciously experience the social context under anesthesia^3^. Findings from Lu and colleagues therefore likely extend to psilocybin what we had previously described for MDMA and ketamine (again, other protocol alterations notwithstanding).

## Discussion and Conclusions

While our initial discovery of psychedelic mediated critical period reopening was made for the social reward learning critical period, more recent evidence demonstrates broader generalizability of psychedelic-mediated reopening to other critical periods. For example, visual critical periods such as those for ocular dominance plasticity (ODP) and orientation selectivity have been reopened by administration of ketamine^9-11^, ibogaine^12^, and LSD^13^ across independently funded studies in multiple labs. Furthermore, six independently funded studies have reproduced our mechanistic findings^2,3^ that metaplasticity and extracellular matrix (ECM) remodeling underlie psychedelic mediated critical period reopening^11,14-18^. Lu et al., fail to cite this literature, thereby misrepresenting the state of the field in their discussion and conclusions.

Interestingly, although social and visual critical periods are measured in different ways, they share sensitivity to many of the same protocol specifications. For example, rearing conditions such as visual, sensorimotor and social complexity significantly impact ODP^19-27^ and the social CPP assay is likewise sensitive to rearing history, such that social reward learning is most robust for same-sex, littermates, housed in groups of 2-5 animals^8,28,29^. Similarly, both sCPP and ODP are sensitive to genetic strain and animal age^29-33^. Additionally, while visual critical period assays are sensitive to luminosity, orientation, spatial frequency of visual gratings and whether the animal is freely moving or under static conditions^34-36^, social CPP is sensitive to the texture and smell of bedding cues, and whether the animal can use them for nesting or burrowing^4,8^. Finally, like psychedelic mediated reopening of the social reward learning critical period, deprivation induced reopening of the ODP critical period is context dependent^19,20,37^.

Taken together, the current analysis challenges the anthropocentric view that mechanistic studies in mice should utilize only simple behavioral assays that are resilient to context dependent effects. While Lu and colleagues imply that strict protocol adherence requirements signify a lack of robustness, our comparative analysis reveals that the exquisite sensitivity of the social CPP assay enables recapitulation of psychedelics’ most behaviorally and therapeutically salient properties, namely their dose, learning, and context dependence^38^. Moreover, alterations to these parameters in Lu et al., *Nature Neuroscience*, 2026 are neither slight, nor insignificant; thus, these protocol modifications falsify ‘failure to replicate’ claims made therein.

## Methods

Social conditioned place preference methodologies and data sets were retrieved across three publications (Nardou et al., 2019; Nardou et al., 2023; and Lu et al., 2026). Protocols were screened for dosage of drug used, context paired with psychedelic administration, duration of pre- and post-test trials, duration and frequency of training sessions, sex, age and strain of animals. While Nardou et al., 2019 used both sexes for establishing the critical period across development, this behavior was not sexually dimorphic, so subsequent studies including those reported in Nardou et al., 2023 used male mice. Lu et al., 2026 used pooled data from both males and females, which were both included in our re-analysis. The Lu et al., 2026 raw data set is reported as percentage of time spent in both the pre- and the post-test sessions. To maintain a consistent reference point, we assumed a 30 min test and converted the percentages of time spent on the pre- and post-test social bedding side vs the pre-test social bedding side to raw numbers (s). Average times were calculated for the pre- and post-test and presented as mean ± sem. Two-tailed paired *t*-tests were computed using statistical analysis software GraphPad Prism to compare pre-vs post-test values.

## CONTRIBUTIONS

GD AND MW wrote and edited the manuscript. MW reanalyzed published data and made the figure.

## ACKNOWLEDGEMENTS

We thank members of the Dölen laboratory for helpful comments. We are also grateful for research funding from the Bob and Renee Parsons Foundation (GD); The Steve and Alex Cohen Foundation (GD), The Silicon Valley Community Foundation (GD), The Alison Family Foundation (GD), and the National Institutes of Health: R01MH117127 (GD), R01HD098184 (GD)

